# 10 Year Epidemiologic data of Parkinson’s Disease: A Nationwide Population-based Retrospective Cohort of South Korea

**DOI:** 10.1101/253682

**Authors:** Hyoung Seop Kim, Joon-Byum Kwon, Hyun-Sun Lim, Jiook Cha, Hye Won Kim

**Affiliations:** Department of Physical Medicine and Rehabilitation, National Health Insurance Service Ilsan Hospital; Department and Research Institute of Rehabilitation Medicine, Yonsei University College of Medicine; Research and Analysis Team, National Health Insurance Service Ilsan Hospital; Department of Psychiatry, New York State Psychiatric Institute, Columbia University Medical Center; Rehabilitation Medicine, Bucheon St. Mary’s Hospital, The Catholic University of Korea, Bucheon, Republic of Korea

**Author notes:** **Corresponding author: Name: Hye Won Kim M.D., Ph.D**. Address: Department of Rehabilitation Medicine, Bucheon St. Mary’s Hospital, 327 Sosa-ro, Wonmi-gu, Bucheon, Gyeonggi-do, Republic of Korea; Zip Code: 14647, Country: Republic of Korea.

**Keywords:** Cause of Death, Incidence, Parkinson’s Disease, Prevalence, Survival Rate

## Abstract

**Objectives:** The aims of this study were to determine the prevalence, incidence, and P/I ratio of Parkinson’s disease (PD) in South Korea and to present basic epidemiological information on PD patients for making effective health policies.

**Methods:** We used National Health Insurance Service-National Sample Cohort (KNHIS-NSC) data to analyze the prevalence, incidence, and P/I ratio of PD from 2003 to 2013 and then followed up using the NHID in 2008 to obtain the hazard ratio (HR) of death in PD itself and other comorbidities from 2008 to 2013.

**Results:** The prevalence and incidence of PD increased rapidly from 72.9 and 32.8 in 2003 to 213.4 and 58.0 in 2013, and the P/I ratio increased from 2.22 in 2003 to 3.62 in 2013. The prevalence, incidence, and P/I ratio of PD were all higher in women than in men. The hazard ratio for death was significantly higher in PD patients (15.36) compared to subjects without the disease. Stroke was the most frequent cause of death in the PD patient population followed by cancer and pneumonia.

**Conclusion:** The prevalence, incidence, and P/I ratio of PD rapidly increased as the years progressed. This indirectly proves that the health insurance system in Korea is efficient and has allowed patients with PD to access medical facilities more easily. However, a newer public healthy strategy should be established for patients with PD because PD itself has a high HR for death, and patients with PD have a high mortality rate when stroke and pneumonia are also involved.

**Disclosure:** All authors have reported no biomedical interests and potential conflicts of interests.

## Introduction

Korea is moving toward an aged society with over 14% of the population aged 65 and older, which is higher than the global percentage of over 7% of the population aged 65 and older, along with experiencing rapid social and economic progress. In 1980, the population of Koreans aged 65 and older was only 3.8%, but increased to 13.0% in 2015 and is expected to surge to 35.9% by 2050.^1^ According to the report on “Global Health and Aging” by the NIH in 2015, South Korea is expected to have the second highest proportion of the population aged 65 or older in the world by 2050 at 35.9% (15,570,000/43,370,000 people), behind Japan at 40.1%.^2^ Among the developed countries in the Western society, France took 157 years to become a super-aged society (with over 217% of the population aged 65 and older) from an aging society, while England took 100 years and the United States took 89 years. In contrast, the rate at which the Asian countries are expected to become a super-aged society is much higher than Western countries: China (34 years), Thailand (35 years), Japan (37 years), and South Korea with the fastest rate among all Asian countries (27 years).

Parkinson’s disease (PD) is a typical degenerative brain disease with four cardinal motor symptoms including bradykinesia, tremor, rigidity, and gait disturbance as well as various non-motor symptoms.^3^ As the disease progresses, it causes loss of independence of gait and daily activities resulting in a bed-ridden state and even death. Due to these problems, PD has been one of the major health problems causing social and economic burden in developed countries.^4^ Although the prevalence and incidence of the disease may vary depending on the country and study, the general trend shows an increase in the prevalence and incidence as the population ages.^3, 5-11^ This phenomenon is believed to further increase as the average life expectancy increases.

Until recently, there has been no nationwide epidemiologic study on patients with PD in South Korea. Therefore, we used the database from the National Health Insurance Service system to analyze changes in the prevalence, incidence, P/I ratio, major causes of death, and hazard ratio (HR) for death in PD patients. The main aim of this study was to determine the prevalence, incidence, and P/I ratio of PD in South Korea. These results may help accurately assess the efficiency of the current geriatric health care policy in South Korea, and provide information to aid in the development of a new policy for PD patients.

## Materials and Methods

### Data Sources

The Korean National Health Insurance Service Corporation (KNHIS) has been distributing the National Health Information Database (NHID) of the National Health Insurance Service in South Korea since 2015. The NHID sample cohort is a nationally representative random data group, which consists of 2.2% of the total number of Koreans with health insurance. Since 2002, it has accumulated more data every year. To extract data on patients with PD, we used the Korean Classification of Diseases (KCD) as disease classification codes, which were modified from the International Classification of Diseases (ICD). Patients diagnosed with PD and primary Parkinsonism with the code G20 were included. Patients with the codes for secondary Parkinsonism (G21) or Parkinsonism in other diseases (G22) were excluded.

We used the NHID sample cohort to analyze the prevalence, incidence, and P/I ratio of PD according to gender and age (under 50, 50-59, 60-69, 70-79, and 80 and over) from 2003 to 2013.

In addition, we used the NHID cohort in 2008 to determine the 5-year survival rate for the main causes of death and the hazard ratios of PD itself and comorbidities from 2008 to 2013 (Fig. 1). The major causes of death were compared in PD patients and subjects without PD, while the hazard ratios were obtained by comparison to the control population without PD.

**Fig. 1.**
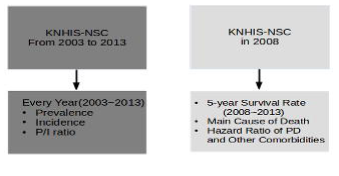
Schematic Diagram of the Study Abbreviation: KNHIS-NSC, Korean National Health Insurance Service-National Sample Cohort

## Statistics

The statistical package SAS for Windows, version 9.4 (SAS Inc, Cary, NC, USA) was used to analyze the data. P < 0.05 was defined as statistically significant. We used the Breslow-Day test to verify yearly changes in the prevalence, incidence, and P/I ratio. We used multiple Cox proportional hazard regression models to obtain the hazard ratios of PD itself and comorbidities. The survival curve estimate in subjects with and without PD was obtained using the Kaplan-Meier method.

## Results

Table 1 shows the prevalence and the incidence of PD according to age and gender from 2003 to 2013. The prevalence and incidence of PD in all subjects changed from 72.9 and 32.8 in 2003 to 213.4 and 58.0 in 2013. As the years progressed, the prevalence and incidence tended to increase rapidly. When comparing the prevalence and incidence of PD between the male and female populations, females had both higher prevalence and incidence each year. In terms of the age distribution of the prevalence and incidence of PD, individuals in the 70s and in the 80s and older were the dominant populations every year. In the male population, the two indexes have been larger in the population in the 80s and older than the population in the 70s since 2003, but in the female population, the prevalence in the 70s was larger until 2010 and the incidence in the 70s was larger until 2008.

**Table 1.** The prevalence and incidence of Parkinson’s disease from 2003 to 2013

| Year | Sex | Male |  |  |  |  |  | Female |  |  |  |  |  | All subjects |
| --- | --- | --- | --- | --- | --- | --- | --- | --- | --- | --- | --- | --- | --- | --- |
|  | Age | <50 | 50-59 | 60-69 | 70-79 | >80 | Sum | <50 | 50-59 | 60-69 | 70-79 | >80 | Sum | Total |
| 2003 | Population | 405,663 | 49,385 | 35,786 | 14,063 | 4,315 | 509,212 | 382,824 | 49,594 | 42,005 | 23,546 | 10,287 | 508,256 | 1,017,468 |
|  | Prevalence | 7.9 | 99.2 | 265.5 | 675.5 | 950.2 | 61.3 | 9.7 | 92.8 | 292.8 | 764.5 | 427.7 | 84.6 | 72.9 |
|  | Incidence | 4.4 | 46.6 | 109.0 | 312.9 | 417.2 | 27.9 | 6.3 | 38.3 | 114.3 | 348.3 | 184.7 | 37.8 | 32.8 |
| 2004 | Population | 400,698 | 51,647 | 36,236 | 15,157 | 4,485 | 508,223 | 379,016 | 51,398 | 42,337 | 24,975 | 10,631 | 508,357 | 1,016,580 |
|  | Prevalence | 6.2 | 87.1 | 300.8 | 673.0 | 914.2 | 63.4 | 6.3 | 89.5 | 330.7 | 732.7 | 536.2 | 88.5 | 75.9 |
|  | Incidence | 3.7 | 27.1 | 127.0 | 257.3 | 289.9 | 25.0 | 3.4 | 27.2 | 146.4 | 252.3 | 216.4 | 34.4 | 29.7 |
| 2005 | Population | 395,794 | 55,103 | 36,317 | 16,405 | 4,698 | 508,317 | 373,779 | 55,247 | 41,919 | 26,350 | 11,208 | 508,503 | 1,016,820 |
|  | Prevalence | 4.3 | 78.0 | 305.6 | 822.9 | 872.7 | 68.3 | 5.6 | 76.0 | 350.7 | 770.4 | 660.2 | 95.8 | 82.0 |
|  | Incidence | 1.5 | 23.6 | 101.9 | 317.0 | 276.7 | 23.8 | 2.9 | 23.5 | 116.9 | 254.3 | 187.4 | 31.7 | 27.7 |
| 2006 | Population | 384,761 | 57,359 | 36,439 | 17,364 | 4,885 | 500,808 | 363,267 | 57,363 | 41,487 | 27,520 | 11,560 | 501,197 | 1,002,005 |
|  | Prevalence | 5.5 | 71.5 | 318.3 | 921.5 | 1207.8 | 79.3 | 8.5 | 81.9 | 392.9 | 879.4 | 743.9 | 113.5 | 96.4 |
|  | Incidence | 2.6 | 24.4 | 107.0 | 322.5 | 429.9 | 28.0 | 5.8 | 34.9 | 149.4 | 261.6 | 285.5 | 41.5 | 34.7 |
| 2007 | Population | 386,671 | 60,989 | 38,245 | 18,897 | 5,207 | 510,009 | 365,380 | 60,776 | 43,157 | 28,999 | 12,422 | 510,734 | 1,020,743 |
|  | Prevalence | 6.2 | 82.0 | 316.4 | 910.2 | 1440.4 | 86.7 | 6.0 | 79.0 | 433.3 | 1079.4 | 845.3 | 132.2 | 109.4 |
|  | Incidence | 3.1 | 27.9 | 99.4 | 275.2 | 480.1 | 28.2 | 1.9 | 19.7 | 155.3 | 362.1 | 330.1 | 45.4 | 36.8 |
| 2008 | Population | 374,855 | 62,553 | 38,594 | 19,669 | 5,348 | 501,019 | 353,093 | 61,857 | 42,949 | 29,105 | 12,762 | 499,766 | 1,000,785 |
|  | Prevalence | 7.5 | 113.5 | 321.3 | 1001.6 | 1720.3 | 102.2 | 9.4 | 101.9 | 470.3 | 1254.1 | 1042.2 | 159.3 | 130.7 |
|  | Incidence | 3.7 | 46.4 | 90.7 | 396.6 | 673.2 | 38.3 | 5.1 | 38.8 | 174.6 | 453.5 | 438.8 | 61.0 | 49.7 |
| 2009 | Population | 367,753 | 65,925 | 39,278 | 20,990 | 5,743 | 499,689 | 346,395 | 65,161 | 43,263 | 30,361 | 13,658 | 498,838 | 998,527 |
|  | Prevalence | 8.2 | 101.6 | 328.4 | 1110.1 | 1880.6 | 113.5 | 8.1 | 102.8 | 483.1 | 1347.1 | 1339.9 | 179.6 | 146.5 |
|  | Incidence | 4.4 | 27.3 | 86.6 | 376.4 | 592.0 | 36.2 | 3.8 | 38.4 | 171.1 | 418.3 | 512.5 | 61.9 | 49.1 |
| 2010 | Population | 362,215 | 70,213 | 40,443 | 22,250 | 6,217 | 501,338 | 340,731 | 69,737 | 43,917 | 31,619 | 14,689 | 500,693 | 1,002,031 |
|  | Prevalence | 6.6 | 79.8 | 341.2 | 1020.2 | 1898.0 | 112.3 | 8.2 | 98.9 | 423.5 | 1280.9 | 1259.5 | 174.4 | 143.3 |
|  | Incidence | 2.5 | 21.4 | 98.9 | 346.1 | 611.2 | 35.7 | 4.4 | 37.3 | 107.0 | 322.6 | 388.1 | 49.3 | 42.5 |
| 2011 | Population | 357,459 | 74,945 | 40,625 | 23,808 | 6,591 | 503,428 | 335,841 | 74,588 | 43,828 | 33,188 | 15,608 | 503,053 | 1,006,481 |
|  | Prevalence | 8.7 | 97.4 | 356.9 | 1201.3 | 2169.6 | 134.7 | 11.6 | 100.6 | 476.9 | 1503.6 | 1941.3 | 223.6 | 179.2 |
|  | Incidence | 4.8 | 25.4 | 115.7 | 336.0 | 576.5 | 39.9 | 6.3 | 25.5 | 139.2 | 439.9 | 602.3 | 67.8 | 53.9 |
| 2012 | Population | 353,134 | 77,670 | 41,903 | 25,800 | 7,107 | 505,614 | 331,811 | 77,045 | 44,755 | 35,086 | 16,812 | 505,509 | 1,011,123 |
|  | Prevalence | 8.5 | 91.4 | 367.5 | 1240.3 | 2237.2 | 145.2 | 9.6 | 105.1 | 480.4 | 1604.6 | 2123.5 | 246.9 | 196.0 |
|  | Incidence | 3.7 | 25.8 | 107.4 | 329.5 | 492.5 | 39.2 | 3.3 | 29.9 | 127.4 | 430.4 | 499.6 | 64.5 | 51.8 |
| 2013 | Population | 349,069 | 80,399 | 43,260 | 26,939 | 7,622 | 507,289 | 328,242 | 79,084 | 46,076 | 36,137 | 17,902 | 507,441 | 1,014,730 |
|  | Prevalence | 11.2 | 90.8 | 409.2 | 1280.7 | 2348.5 | 160.3 | 11.3 | 113.8 | 462.3 | 1732.3 | 2161.8 | 266.6 | 213.4 |
|  | Incidence | 6.3 | 27.4 | 131.8 | 378.6 | 564.2 | 48.5 | 6.1 | 41.7 | 108.5 | 448.3 | 435.7 | 67.6 | 58.0 |

The P/I ratio has increased with age for 11 years from 2003 to 2013 (Table 2). The ratio in all subjects changed from 2.22 in 2003 to 3.62 in 2013, which was statistically significant (p<0.0001). There has also been a gradual increase in the yearly change in the ratio in both the male and female population with statistical significance (p=0.0036 and p<0.0001, respectively) from 2003 to 2013. When comparing the ratio between the male and female population, the female population had a higher P/I ratio every year. We also compared the yearly change in P/I ratio in each age group by decade and found that the female group in the 70s and in the 80s and older showed a gradual increase in the P/I ratio with statistical significance (p=0.0108 and p<0.001) as the year progressed, while the male group did not show an increase in any age group.

**Table 2.** The prevalence/incidence ratio of Parkinson’s disease from 2003 to 2013

| Year | Sex | Male |  |  |  |  |  | Female |  |  |  |  |  | All subjects |
| --- | --- | --- | --- | --- | --- | --- | --- | --- | --- | --- | --- | --- | --- | --- |
|  | Age | <50 | 50-59 | 60-69 | 70-79 | >80 | Sum | <50 | 50-59 | 60-69 | 70-79 | >80 | Sum | Total |
| 2003 |  | 1.78 | 2.13 | 2.44 | 2.16 | 2.28 | 2.20 | 1.54 | 2.42 | 2.56 | 2.20 | 2.32 | 2.24 | 2.22 |
| 2004 |  | 1.67 | 3.24 | 2.37 | 2.62 | 3.15 | 2.54 | 1.85 | 3.29 | 2.26 | 2.91 | 2.48 | 2.58 | 2.56 |
| 2005 |  | 2.83 | 3.31 | 3.00 | 2.60 | 3.15 | 2.87 | 1.91 | 3.23 | 3.00 | 3.03 | 3.52 | 3.02 | 2.95 |
| 2006 |  | 2.10 | 2.93 | 2.98 | 2.86 | 2.81 | 2.84 | 1.47 | 2.35 | 2.63 | 3.36 | 2.61 | 2.73 | 2.79 |
| 2007 |  | 2.00 | 2.94 | 3.18 | 3.31 | 3.00 | 3.07 | 3.14 | 4.00 | 2.79 | 2.98 | 2.56 | 2.91 | 2.99 |
| 2008 |  | 2.00 | 2.45 | 3.54 | 2.52 | 2.56 | 2.67 | 1.83 | 2.63 | 2.69 | 2.76 | 2.38 | 2.61 | 2.64 |
| 2009 |  | 1.88 | 3.72 | 3.79 | 2.95 | 3.18 | 3.14 | 2.15 | 2.68 | 2.82 | 3.22 | 2.61 | 2.90 | 3.02 |
| 2010 |  | 2.67 | 3.73 | 3.45 | 2.95 | 3.11 | 3.15 | 1.87 | 2.657 | 3.96 | 3.97 | 3.25 | 3.53 | 3.34 |
| 2011 |  | 1.82 | 3.84 | 3.09 | 3.58 | 3.76 | 3.38 | 1.86 | 3.95 | 3.43 | 3.43 | 3.22 | 3.3 | 3.34 |
| 2012 |  | 2.30 | 3.55 | 3.42 | 3.76 | 4.54 | 3.71 | 2.91 | 3.52 | 3.771 | 3.73 | 4.25 | 3.83 | 3.77 |
| 2013 |  | 1.77 | 3.32 | 3.11 | 3.38 | 4.16 | 3.30 | 1.85 | 2.73 | 4.26 | 3.86 | 4.96 | 3.94 | 3.62 |
|  | P-value | 0.9956 | 0.858 | 0.7832 | 0.1992 | 0.4554 | 0.0036 | 0.9216 | 0.892 | 0.066 | 0.0108 | 0.001 | <0.0001 | <0.0001 |

Unit: %

The results of the major causes of death and hazard ratios of death based on the analysis from 2008 to 2013 are shown in Table 3. The major causes of death in the total population were cancer (6,943 deaths/24,559 total deaths; 28.27%), stroke (2,221 deaths; 9.04%), ischemic heart disease (1,253 deaths; 5.10%), pneumonia (805 deaths; 3.28%), and other heart disease. On the other hand, stroke was the most frequent cause of death in patients with PD (51 deaths/408 total deaths; 12.5%), followed by cancer (44 deaths; 10.78%), pneumonia (15 deaths; 3.68%), and ischemic heart disease (14 deaths; 3.43%).

**Table 3.**
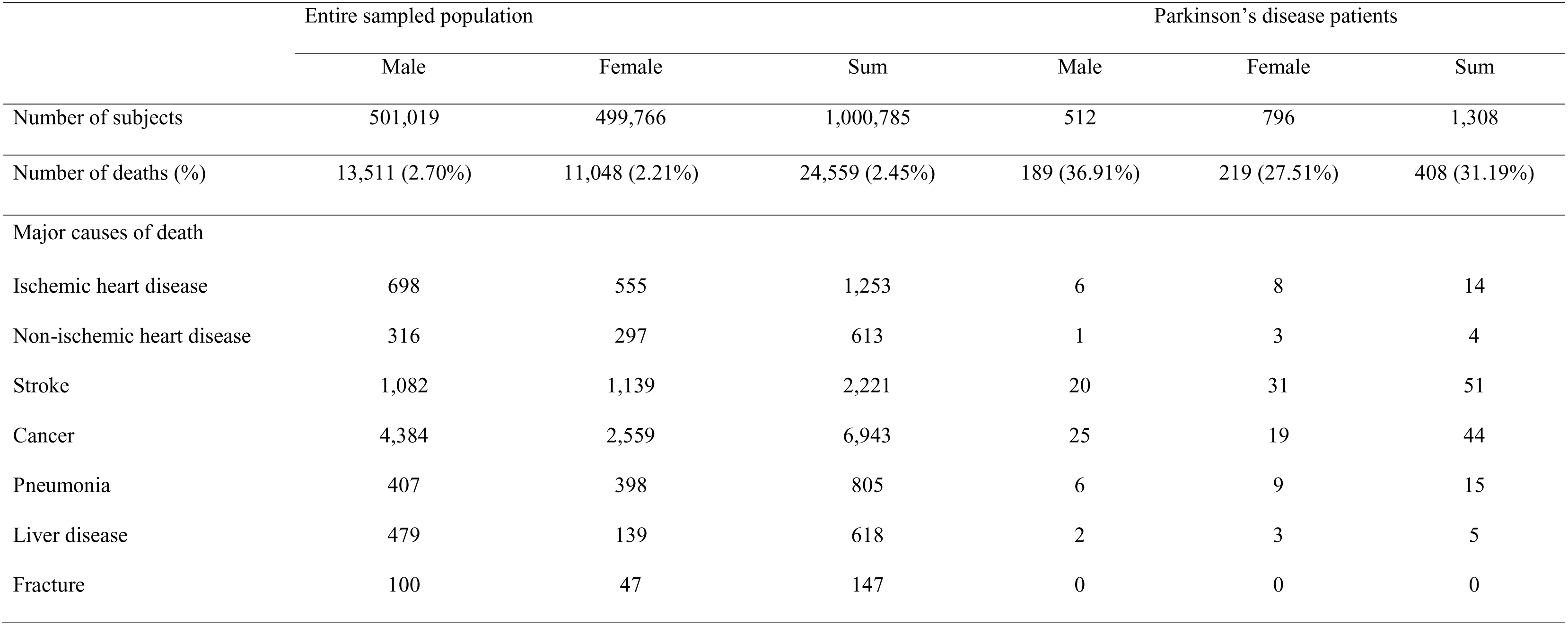
Comparison of causes of death in the entire sampled population and patients with Parkinson’s disease from 2008 to 2013

|  | Entire sampled population |  |  | Parkinson’s disease patients |  |  |
| --- | --- | --- | --- | --- | --- | --- |
|  | Male | Female | Sum | Male | Female | Sum |
| Number of subjects | 501,019 | 499,766 | 1,000,785 | 512 | 796 | 1,308 |
| Number of deaths (%) | 13,511 (2.70%) | 11,048 (2.21%) | 24,559 (2.45%) | 189 (36.91%) | 219 (27.51%) | 408 (31.19%) |
| Major causes of death |  |  |  |  |  |  |
| Ischemic heart disease | 698 | 555 | 1,253 | 6 | 8 | 14 |
| Non-ischemic heart disease | 316 | 297 | 613 | 1 | 3 | 4 |
| Stroke | 1,082 | 1,139 | 2,221 | 20 | 31 | 51 |
| Cancer | 4,384 | 2,559 | 6,943 | 25 | 19 | 44 |
| Pneumonia | 407 | 398 | 805 | 6 | 9 | 15 |
| Liver disease | 479 | 139 | 618 | 2 | 3 | 5 |
| Fracture | 100 | 47 | 147 | 0 | 0 | 0 |

Table 4 shows that the hazard ratio of death was much higher in PD patients compared with subjects without PD (HR: 15.36). The survival rate was 95% or more in subjects without PD, but dropped to less than 70% in patients with PD after 1,500 days since January 1, 2008 (Fig. 2). Analysis of the hazard ratio of comorbidities within the patient population showed that the hazard ratio of stroke was the highest at 18.56, followed by pneumonia at 15.05, ischemic heart disease at 8.99, and non-ischemic heart disease at 5.65.

**Fig. 2.**
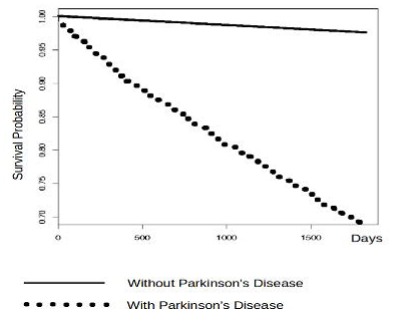
Comparison of the 5-Year Survival Rate between Subjects without PD and Patients with PD

**Table 4.**
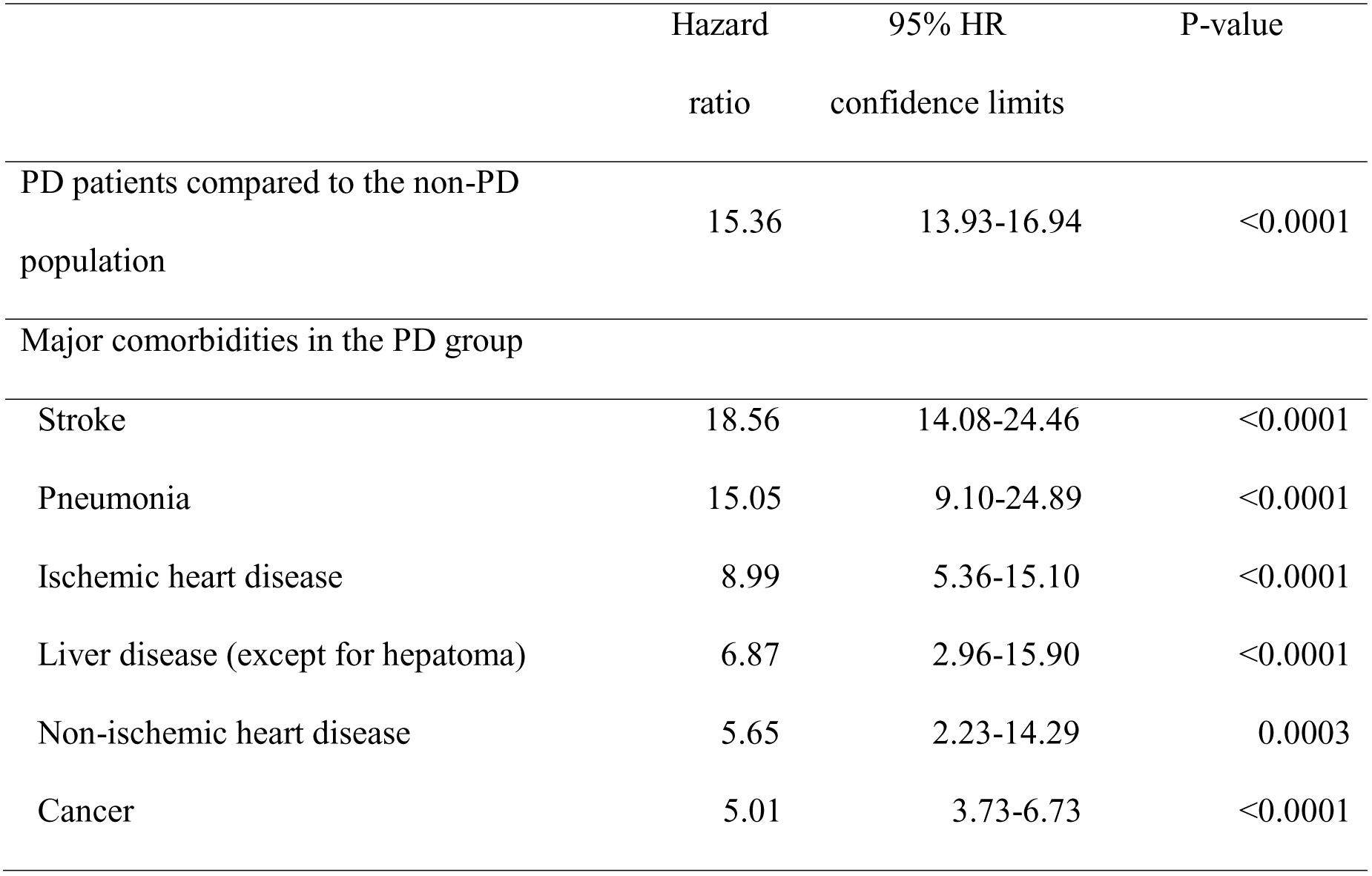
Hazard ratios (HR) of Parkinson’s disease (PD) and major comorbidities in PD patients

|  | Hazard<br>ratio | 95% HR<br>confidence limits | P-value |
| --- | --- | --- | --- |
| PD patients compared to the non-PD<br>population | 15.36 | 13.93-16.94 | <0.0001 |
| Major comorbidities in the PD group |  |  |  |
| Stroke | 18.56 | 14.08-24.46 | <0.0001 |
| Pneumonia | 15.05 | 9.10-24.89 | <0.0001 |
| Ischemic heart disease | 8.99 | 5.36-15.10 | <0.0001 |
| Liver disease (except for hepatoma) | 6.87 | 2.96-15.90 | <0.0001 |
| Non-ischemic heart disease | 5.65 | 2.23-14.29 | 0.0003 |
| Cancer | 5.01 | 3.73-6.73 | <0.0001 |

## Discussion

Studies related to the epidemiology of PD patients in other countries have shown controversial results depending on the sampled population size of the country or community, or the methodology used such as door-to-door surveys versus medical records.3-6, 8, 12-16

Muangpaisan et al. studied the epidemiology of the disease in Asia using a meta-analysis of studies published from 1965 to 2008 and reported that the range in standardized all-age prevalence was 51.3 to 176.9 per 100,000 in door-to-door surveys and 35.8 to 68.3 in medical record-based surveys, and the range of the standardized incidence was 8.7 person-years in door-to-door surveys and 6.7 to 8.3 in record-based surveys.^3^ In non-Asian countries by comparison, the range of the standardized prevalence was 101.0 to 439.4 in door-to-door surveys and 61.4 to 141.1 in record-based surveys, while the range of the standardized incidence was 15.4 to 27.6 in door-to-door surveys and 6.1 to 17.4 in record-based surveys. In our study, the prevalence of PD in all subjects increased every year from 72.9 in 2003 to 213.4 in 2013, and the incidence of PD increased every year from 32.8 in 2003 to 58.0 in 2013.

In terms of the prevalence of PD, our finding of the rapid increase every year is similar to previous reports from Israel, France, Japan, and Taiwan.^8, 15-17^ However, studies from the UK and US reported a stable pattern of PD prevalence.^13, 14^ With regard to the incidence of PD, our study revealed an increase in the incidence every year. However, previous reports in the US, Israel, and France^14, 16, 17^ showed a stable pattern of PD incidence over each year. Also, a decreasing pattern of PD incidence was shown in reports from Taiwan and the UK. ^8, 13^

This rapid increase in the prevalence and incidence of PD in South Korea may be explained by the increase in diagnosis of PD due to the ease of accessibility to hospitals along with low medical expenses in South Korea under the universal National Health Insurance Service and the expansion of the National Health Screening Project that is mandatory for every individual above 40 years old. In addition, the elderly population in Korea has increased due to higher average life expectancy because of improvements in economic status and medical technologies as well as enforcement of the social welfare system the National Long-Term Care Service since 2008.

When analyzing the age distribution of the prevalence and incidence of PD, individuals in their 70s and 80s and over were the dominant population every year. In contrast, some previous studies reported an abrupt decline in the prevalence and incidence of PD in individuals in their 80s.^1,3,6^ An explanation for this difference is the possibility of under-diagnosis of PD in the elderly over 80. One previous study reported that the majority of patients with undiagnosed PD were in their 80s or older, which may be due to limitations in access to hospitals and diagnosis masked by other medical comorbidities.^18^ Four door-to-door surveys revealed under-diagnosis of prevalent idiopathic Parkinsonism at rates ranging from 30% to 71%.^6^ Meanwhile, under-diagnosis is rare in South Korea because of the ease in accessibility to hospitals and the National Health Screening Project. Also, in terms of the pathophysiology of PD, the prevalence and incidence should be increasing as the population becomes older because of the nature of this neurodegenerative disorder that causes degeneration of dopaminergic neurons in the brain.

The ratio of prevalence to incidence (P/I ratio) in PD is a useful tool that reflects the average duration of survival and quality of management for the disease.^3^ A higher P/I ratio indicates that better treatment is given to patients with PD. The P/I ratio in our study linearly increased from 2.22 in 2003 to 3.62 in 2013 as the years progressed. Also, there has been a gradual increase in the yearly change in the ratio in both the male and female population with statistical significance (p=0.0036 and p<0.0001, respectively) for 11 years. This indicates that the quality of management for patients with PD has improved. In fact, the Long-Term Care Insurance Service for the elderly in South Korea was introduced in 2008 in addition to the National Health Care Insurance Service. It covers patients under 65 years old with designated neurodegenerative diseases as well as the elderly over 65 years old who have difficulties in activities of daily living regardless of the disease type. Interestingly, we found that the prevalence, incidence, and P/I ratio of PD were higher in women than men. Several studies conducted in Japan on the epidemiology of PD also showed a higher prevalence and incidence of PD in women than men in all age groups.^12, 18-21^ This may be related to the ethnical similarity between Koreans and the Japanese. A possible biological hypothesis to explain this result is that female patients with PD have a more benign phenotype. Additionally, symptoms of PD may develop slower in women because of higher striate dopamine levels, possibly related to the neuroprotective activity of estrogen.^8, 14-16^ On the other hand, a few studies have reported that the prevalence and incidence of PD were higher in men. These studies were mainly conducted in Western Caucasian populations in the US, Australia, Europe, and Finland.^12, 14, 16^ Furthermore, a study in Taiwan found a different result from ours despite a biologically similar study population, using a similar methodology, and operating under a national health insurance system like South Korea.^8^ These different results may be due to demographic characteristics, occupational and environmental differences, and genetic factors.^18^

In the analysis of major causes of death in PD patients, stroke was first, cancer second, pneumonia third, and ischemic heart disease fourth (Table 3). In contrast, the major causes of death in the entire sampled population were cancer first, followed by cerebral stroke second, ischemic heart disease third, and pneumonia fourth. In addition, the hazard ratio of PD itself to death was 15.36, indicating that the survival rate of patients with PD in our study was significantly lower than in subjects without PD (Fig. 2). A similar result was obtained from a study in Taiwan using a door-to-door survey method.^1^ They showed that the relative risk of death in PD patients versus non-PD was 3.38, and the fatality rate for PD after a 7-year follow-up was 40.4%. This previous study also found that the survival rate of patients older than 65 years was significantly lower than that of patients aged 40 to 65 years.^1^ Another study on a European population also reported an increased mortality rate in patients with PD over 70 years old.^3^

In terms of the hazard ratio of comorbidities leading to death in PD patients (Table 4), the hazard ratio of stroke was the highest at 18.56, followed by pneumonia at 15.05, ischemic heart disease at 8.99, and non-ischemic heart disease at 5.65, indicating that the comorbidities of stroke or pneumonia in PD patients had the greatest influence on death.

There are several possible explanations for why stroke was the leading cause of death in PD patients. Patients with PD are more likely to experience a decline in physical activity as the disease progresses, resulting in a bedridden state at the terminal stage. This reduced physical activity and mobility have a negative influence on blood coagulation and viscosity, which increases the chances of cerebral stroke. In the late stage of PD, dysphagia commonly occurs because of impairments in the protection mechanism against aspiration, resulting in silent aspiration of the lungs. This inevitably increases the chances of aspiration pneumonia in PD patients. Moreover, dysphagia and aspiration are exacerbated when stroke also occurs in PD patients.

There has not yet been any report on the hazard ratios of PD itself and comorbidities in PD patients. Therefore, this report is valuable for its contribution to understanding of the characteristics of comorbidities leading to death in patients with PD.

We were also interested in the rate of fractures from falling, which is common among PD patients due to motor symptoms and postural instability^22^, but there was no difference in mortality from falls and fractures between the PD patient population and the population without PD. An assumption can be made that this phenomenon is due to most end-stage PD patients being bed-ridden, remarkably decreasing the risk of falls and fractures.

Since this was a retrospective cohort study, there was a limitation in this research because the diagnosis of PD was only determined by the KCD code, making it impossible to apply the same diagnostic criteria as a prospective study.

The strong points of our study are as follows. Our study was the first in Korea to report the prevalence and incidence, P/I ratio, major causes of death, and hazard ratio of death in PD patients utilizing a nationwide health insurance database. We were able to include a large population for our study while previous studies only included door-to-door surveys or community-based surveys with a limited number of subjects. The National Health Insurance database in South Korea is reliable because the Health Insurance Review and Assessment Service (HIRAS) reviews and supervises the electronic medical records that are submitted from medical facilities. Therefore, the results of this study can be useful for comparing the epidemiology of PD in Korea to the disease in other countries.

## Conclusion

By examining data from the National Health Insurance Service-National Sample Cohort (NHIS-NSC) database from 2003 to 2013, we found that the annual prevalence, incidence, and P/I ratio of PD increased as the years progressed and were higher in women than men. This indirectly proves that the health insurance or welfare system in Korea is efficient and has allowed patients with PD to more easily utilize medical facilities or public services. Also, from the NHIS-NSC data from 2008 to 2013, we found that the leading cause of death in patients with PD was stroke, while cancer was the leading cause in the entire sampled population. The hazard ratio of death in PD patients was very high at 15.36. Stroke and pneumonia were the two major comorbidities causing death in PD patients. This is a somewhat different pattern from that of subjects without PD, suggesting that a new public health strategy should be established for patients with PD.

We hope that the results of this study will be useful for comparing epidemiologic data with other countries and verifying the efficiency of the current geriatric health care service in Korea, along with establishing future health and welfare policies for elderly patients with PD.

